# Immediate visualization of recombination events and chromosome segregation defects in fission yeast meiosis

**DOI:** 10.1101/458398

**Authors:** Dmitriy Li, Marianne Rocl, Raif Yuecel, Alexander Lorenz

**Affiliations:** Institute of Medical Sciences (IMS) and; Iain Fraser Cytometry Centre (IFCC), University of Aberdeen, Foresterhill, Aberdeen AB25 2ZD, UK; Present address: Laboratoire de Biologie du Développement de Villefranche-sur-Mer (LBDV), Sorbonne Université, 06230 Villefranche-sur-Mer, France

**Keywords:** *Schizosaccharomyces pombe*, chromosome segregation, meiotic recombination, spore-autonomous promoters, imaging flow cytometry

## Abstract

*Schizosaccharomyces pombe*, also known as fission yeast, is an established model for studying chromosome biological processes. Over the years research employing fission yeast has made important contributions to our knowledge about chromosome segregation during meiosis, as well as meiotic recombination and its regulation. Quantification of meiotic recombination frequency is not a straightforward undertaking, either requiring viable progeny for a genetic plating assay, or relying on laborious Southern blot analysis of recombination intermediates. Neither of these methods lends itself to high-throughput screens to identify novel meiotic factors. Here, we establish visual assays novel to *Sz. pombe* for characterizing chromosome segregation and meiotic recombination phenotypes. Genes expressing red, yellow, and/or cyan fluorophores from spore-autonomous promoters have been integrated into the fission yeast genomes, either close to the centromere of chromosome I to monitor chromosome segregation, or on the arm of chromosome III to form a genetic interval at which recombination frequency can be determined. The visual recombination assay allows straightforward and immediate assessment of the genetic outcome of a single meiosis by epi-fluorescence microscopy without requiring tetrad dissection. We also demonstrate that the recombination frequency analysis can be automatized by utilizing imaging flow cytometry to enable high-throughput screens. These assays have several advantages over traditional methods for analysing meiotic phenotypes.

## Introduction

Meiosis is a highly-conserved process that produces haploid sex cells (gametes) as an integral part of sexual reproduction (Hunter 2015). During meiosis chromosomes are deliberately broken to initiate homologous (meiotic) recombination that physically connects the equivalent maternal and paternal (homologous) chromosomes, this is absolutely essential for correct chromosome segregation (Petronczki et al. 2003; Lam and Keeney 2015). Only if these connections (chiasmata) are achieved accurately, healthy gametes containing a single chromosome complement will result from the two meiotic cell divisions. In the process homologous chromosomes are re-shuffled and genes are re-assorted; this process provides the genetic diversity that makes individuals unique. Failure to perform meiosis correctly has been shown to cause infertility, miscarriages, and hereditary disorders in mammals (Hassold and Hunt 2001); meiosis is thus fundamental to sexual reproduction.

Meiotic recombination is initiated by Spo11, a TopVI-like transesterase, creating meiotic double-stranded DNA breaks (DSBs) (Lam and Keeney 2015). These DSBs are subsequently repaired by homology-directed repair mechanisms driven by the RecA-family recombinases Rad51 and Dmc1. Rad51 and its meiosis-specific paralogue Dmc1 are supported by a host of ancillary factors through loading Rad51 and or Dmc1 onto a processed DSB site and stabilizing them as multimeric nucleoprotein filaments. These ancillary factors include Rad51-paralogues (Gasior et al. 1998; Grishchuk and Kohli 2003; Bleuyard et al. 2005; Sasanuma et al. 2013; Brown and Bishop 2014; Lorenz et al. 2014; Abreu et al. 2018), and factors evolutionarily unrelated to RecA, such as Rad52, Swi5-Sfr1 & Hop2-Mnd1 (Gasior et al. 1998; Chen et al. 2004; Ellermeier et al. 2004; Zierhut et al. 2004; Petukhova et al. 2005; Kerzendorfer et al. 2006; Vignard et al. 2007; Octobre et al. 2008). In *Sz. pombe* the Hop2-Mnd1 orthologues are called Meu13-Mcp7, and similar to the situation in other eukaryotes meiotic recombination is strongly reduced in their absence (Nabeshima et al. 2001; Saito et al. 2004). Homology-directed repair can follow several pathways, and ultimately results in crossover (CO) and non-crossover recombination outcomes (Phadnis et al. 2011; Hunter 2015). Only COs between homologous chromosomes support the formation of chiasmata, which together with sister chromatid cohesion are needed for proper chromosome segregation (Marston 2014). Cohesion is achieved by the cohesin complex which physically entraps the sister chromatids right after their replication during S phase (Nasmyth and Haering 2009). Cohesin holds sister chromatids together until all chromosomes are properly attached to microtubules in metaphase, at which point the kleisin subunit of cohesin is destroyed and anaphase ensues (Nasmyth and Haering 2009; Marston 2014). To reduce the diploid chromosome complement to a haploid one, meiosis consists of two cell divisions following a single round of DNA replication; special modifications to sister chromatid cohesion have to be in place to enable this. During meiosis I homologous chromosomes are segregated from each other, cohesins are only removed from the chromosome arms, whereas cohesins at centromeres remain protected for the second meiotic division. During meiosis II centromeric cohesin protection is removed to allow sister chromatids to be segregated from each other (Petronczki et al. 2003; Marston 2014). A key centromeric protector is the Mei-S homologue Shugoshin, Sgo (Katis et al. 2004; Kitajima et al. 2004; Marston et al. 2004; Rabitsch et al. 2004), and the absence of Sgo and chiasmata, indeed, generates a strong chromosome segregation defect during meiosis (Hirose et al. 2011).

Here, we establish and characterize visual assays to quantify chromosome segregation defects and meiotic recombination frequency which are new to *Sz. pombe*. Visual assays for determining meiotic recombination frequencies were originally established in *Arabidopsis*, and more recently adapted for budding yeast (Francis et al. 2007; Thacker et al. 2011). These visual recombination assays utilize genes encoding red, yellow, and cyan fluorophores driven by gamete-specific promoters, and are integrated at specific loci on a given chromosome to form genetic intervals. The four products (gametes) of a single meiosis will fluoresce in a color corresponding to the fluorophore gene(s) they receive. In *Arabidopsis*, the fluorophores are expressed from the pollen-specific post-meiotic *LAT52*-promoter, and various genetic intervals (fluorescent-tagged lines, FTLs) have been generated and adopted widely (e.g., Yelina et al. 2013; Séguéla-Arnaud et al. 2017; Kurzbauer et al. 2018). Also, the budding yeast version of the visual recombination assay starts to enjoy popularity and several recent studies used spore-autonomous fluorophore expression to determine meiotic recombination frequency (e.g., Vincenten et al. 2015; Arter et al. 2018; González-Arranz et al. 2018). In yeasts this kind of setup allows assessment of the frequency of exchange of flanking markers (COs) and has advantages over traditional methods for studying meiotic recombination – such as using nutritional markers (White and Petes 1994; Smith 2009) or Southern blotting of DNA from meiotic yeast cells (Hyppa and Smith 2009; Oh et al. 2009): (I) spores can be assessed regardless of their viability (ability to form a visible yeast colony), (II) the simplicity of this method will allow its use for high-throughput genetic screens, and (III) achieving large sample sizes is straightforward when using imaging flow cytometry. Additionally, this can also be used as a tool for monitoring chromosome segregation defects, when different fluorophore markers are inserted close to a centromere (Thacker et al. 2011; this study).

These visual assays represent a novel, powerful, and easy-to-use experimental tool for fission yeast allowing straightforward analysis of chromosome segregation and homologous recombination defects during meiosis. They also enable the identification and characterization of complex phenotypes (single and double CO formation) in high-throughput screens via imaging flow cytometry.

## Materials and Methods

### Molecular and microbiological techniques

Plasmids and details of construction are given in Table S1. DNA modifying enzymes (high-fidelity DNA polymerase Q5, Taq DNA polymerase, T DNA ligase, restriction endonucleases) and the NEBuilder HiFi DNA Assembly Master Mix were obtained from New England BioLabs (NEB), Inc. (Ipswich, MA, USA), and the In-fusion HD Cloning kit from Takara Bio, Inc. (Mountain View, CA, USA). Oligonucleotides (Table S2) were supplied by Sigma-Aldrich Co. (St. Louis, MO, USA). All relevant regions of plasmids were verified by DNA sequencing (Source Bioscience plc, Nottingham, UK). Plasmid sequences are available as supporting online material (https://figshare.com/s/8b8aec968952b862523d).

*Escherichia coli* was grown in LB and SOC media, when appropriate media contained μg/ml Ampicillin (Sambrook and Russell 2000). Competent *E. coli* XL1-blue cells (Agilent Technologies, Santa Clara, CA, USA) were transformed following the protocol provided by the manufacturer.

*Schizosaccharomyces pombe* strains (Table S3) were cultured on yeast extract (YE), and on yeast nitrogen base glutamate (YNG) agar plates containing the required supplements (concentration μg/ml on YE, and μg/ml on YNG). Crosses were performed on malt extract (ME) agar with the required amino acids (concentration μg/ml). Fission yeast transformations were performed using a standard Li-acetate protocol (Brown and Lorenz 2016). Spore-autonomously expressed fluorophore genes were targeted to their intended sites using flanking homologous sequences which were provided via various strategies (Table S1) (Bähler et al. 1998; Matsuyama et al. 2004; Gregan et al. 2006). Construction of the *hphMX4*-marked *meu13*Δ-*22* strain UoA585 by marker swap from *meu13*Δ::*ura4*+ has been described elsewhere (Lorenz 2015), the *meu13*Δ-*43*::*natMX4* strain UoA723 was derived by transforming an appropriate marker swap cassette amplified by PCR (oligonucleotides oUA and oUA102, Table S2) from pALo into UoA585 (*meu13*Δ-*22*::*hphMX4*) (Lorenz 2015; Brown and Lorenz 2016). Strains carrying the *meu13*Δ-*22*, *meu13*Δ-*43*, *sgo1*Δ, and *rec12*Δ*-169* alleles were derived by crossing from UoA585, UoA723, JG17888, and GP3717, respectively (Davis and Smith 2003; Gregan et al. 2005; Lorenz 2015).

Spore viability by random spore analysis and meiotic recombination assays have been performed as previously described (Osman et al. 2003; Smith 2009; Sabatinos and Forsburg 2010; Lorenz et al. 2012).

### Microscopy

For microscopy cells from sporulating cultures were suspended in sterile demineralized water, and spotted onto microscopic slides. After placing a cover slip over the cell suspension, cells were immobilized by squashing the slide in a filter paper block, and afterwards the cover slip was sealed with clear nail varnish. Microscopic analysis was done using a Zeiss Axio Imager.M2 (Carl Zeiss AG, Oberkochen, Germany) epi-fluorescence microscope equipped with the appropriate filter sets to detect red, yellow, and cyan fluorescence. Black-and-white images were taken with a Zeiss AxioCam MRm CCD camera controlled by AxioVision 40 software v4.8.2.0. Images were pseudo-colored and overlayed using Adobe Photoshop CC (Adobe Systems Inc., San José, CA, USA). Images of mature 4-spored asci were evaluated manually, data was collected and analyzed in Microsoft Excel MSO (version 16.0.4738.1000, 32-bit). 157

### Imaging flow cytometry

The ImageStreamX Mark II (Merck KGaA, Darmstadt, Germany) is an imaging flow cytometer, where an image of each individual cell is acquired as it flows through the cytometer. It measures hundreds of thousands of individual cells in minutes, combining the high-throughput capabilities of conventional flow cytometry with single-cell imaging. The ImageStream measures not only total fluorescence intensities but also the spatial image of the fluorescence plus bright-field and dark-field images of each cell in a population.

For imaging flow cytometry cellular material containing asci was suspended in 1× PBS, pH 7. (g/l NaCl, g/l KCl, 1. g/l Na_2_HPO_4_·7H_2_O, g/l anhydrous KH_2_PO_4_), harvested by centrifugation (6, ×g, sec), and re-suspended in 1× PBS, pH 7.5. Data was acquired on the ImageStreamX Mark II using INSPIRE acquisition software (Merck kGaA). Cellular parameters were measured in Channel (Brightfield, BF), Channel (GFP*, a yellow-shifted version of green fluorescent protein, using a nm laser), Channel (RFP, red fluorescent protein, nm), Channel (CFP, cyan fluorescent protein, nm), and Channel (side scatter, nm) with magnification set to ×60. Briefly, objects of interest (asci) with a BF ‘Area’ of 50μm^2^ to 200μm^2^ and an ‘Aspect Ratio’ (ratio of minor axis to major axis) lower than 0. (“doublet area”) were selected. Focused cells were identified by a ‘Gradient RMS’ feature value of or higher. A typical file contained about 25, focused yeast cells.

Data evaluation for identification of asci and spore phenotyping were performed using IDEAS software (version 6.2; Merck). A focused population of asci were identified within the “doublet area” and based on the features ‘Modulation’ for fluorescent channels and ‘Intensity’ for side scatter (SSC) using the custom masks ‘Morphology’ and ‘Object(tight)’, respectively. Further refinement was performed each on RFP, GFP* and CFP fluorescence via ‘Intensity’. Following analysis of the merged triple fluorescent population using ‘Length’ and ‘Elongatedness’ features (custom BF mask “AdaptiveErode, M01, Ch01, 75”) resulted in identification of asci of interest. Finally, spore phenotype analysis was conducted by evaluating the fluorescent area using custom masks for each fluorescent intensity (GFP* intensity 200-4095, RFP intensity 75-and CFP intensity 150-4095) and by applying Boolean algebra to identify particular combinations of fluorescent colors. Asci with a mask area larger than 3μm^2^ were considered positive for a particular spore phenotype.

## Results and Discussion

### Identifying spore-autonomous promoters in *Schizosaccharomyces pombe*

To test whether a particular upstream regulatory sequence is a spore-autonomous promoter (Thacker et al. 2011) we cloned a 700-931bp region upstream of the start codon of the *Sz. pombe eis1*, *pil2*, *eng2*, *agn2*, and *mde10* genes in front of a cyan (*mCerulean*) or red (*tdTomato*) fluorophore gene inserted in pDUAL, a vector restoring *leu1*+ by integrating at the *leu1-32* mutant locus (Matsuyama et al. 2004). Fluorophore genes were terminated by *Saccharomyces* spp. *PGK1* downstream regulatory sequence: *TPGK1* from *S. bayanus* for *mCerulean*, and *T_PGK1_* from *S. kudriavzevii* for *tdTomato* (Thacker et al. 2011). The candidate promoters were selected on the basis of its corresponding gene being upregulated during late meiosis or sporulation (Mata et al. 2002): *eng2*, *agn2* & *mde10* code for proteins involved in spore wall formation, *eis1* encodes an eisosome assembly protein, and *pil2* a component of the eisosome. The promoter of *S. cerevisiae YKL050c* has previously been shown to support spore-autonomous expression of fluorophores in budding yeast (Thacker et al. 2011), *Sz. pombe eis1* is the single homologue of the *S. cerevisiae* paralogue pair *EIS1* and *YKL050c*. The resulting plasmids (pALo139, pALo140, pALo141, pALo142, pALo175; Table S1) were digested with *Apa*I to release the *leu1*+ integration cassettes containing the constructs; these were transformed into *h+* and *h-* fission yeast strains (ALP729 and FO652) carrying the *leu1-32* mutation. Two *leu1*+ strains of different mating types carrying differently colored fluorophore constructs were crossed to each other and presence or absence of spore-specific fluorescence was recorded on an epi-fluorescence microscope. *P_eng_2*, *P_agn2_*, and *P_mde10_* failed to produce fluorescence levels visible under the microscope (data not shown). *P_eis1_* and *P_pil2_* were strong spore-autonomous promoters yielding clear red or cyan fluorescence in spores of mature asci (data not shown).

To avoid ectopic recombination events between the *P_eis1_*- and *P_pil2_*-constructs and the upstream regions of endogenous *eis1* and *p_il2_*, we decided to follow a similar strategy as Keeney and co-workers (Thacker et al. 2011), and investigated whether *P_eis1_* and *P_pil2_* from *Schizosaccharomyces* species other than *Sz. pombe* can be used as spore-autonomous promoters in *Sz. pombe*. Indeed, the upstream sequences of the *Sz. japonicus eis1* and *pil2*-homologues *SJAG_04227* and *SJAG_02707,* as well as the regions upstream of *Sz. cryophilus* and *Sz. octosporus pil2*-homologues *SPOG_00147* and *SOCG_04642*, cloned in front of fluorophores produced strong fluorescence in spores of *Sz. pombe* asci (Fig. 1a). *P_SJAG_04227_*, *P_SPOG_00147_*, and *P_SOCG_04642_* were selected to drive tdTomato (red fluorescence protein, from now called RFP), GFP* (yellow-shifted green fluorescence protein, terminated by *T_PGK1_* from *S. mikatae*) (Griesbeck et al. 2001; Thacker et al. 2011), and mCerulean (cyan fluorescence protein, from now on called CFP) expression in all experimental constructs (Fig. 1).

**Figure 1.**
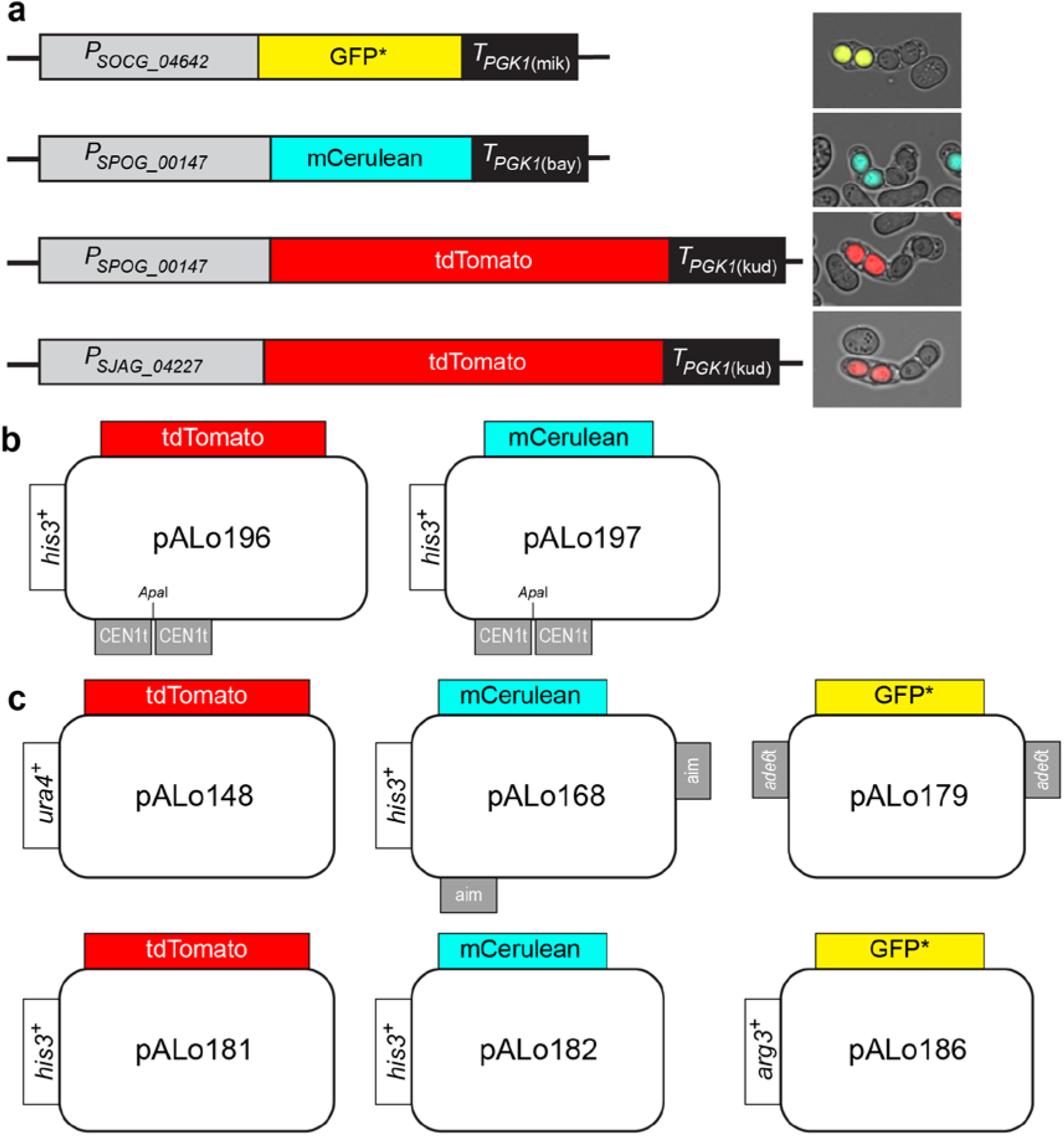
Spore-autonomous expression of fluorophores. (**a**) Schematic and examples of main constructs, *P_SOCG_04642_-GFP*-T_PGK1(mik)_* from strain UoA585694, *P_SPOG_00147_-mCerulean-T_PGK1(bay)_* from strain UoA585727, *P_SPOG_00147_-tdTomato-T_PGK1(kud)_* from strain UoA585726, and *P_SJAG_04227_-tdTomato-T_PGK1(kud)_* from strain UoA585694. (**b**) Plasmid maps of *CEN1*-targeting (CEN1t) constructs using the *Sz. octosporus SPOG_00147* (*pil2*) promoter to drive RFP (tdTomato) and CFP (mCerulean) expression. (**c**) Plasmid maps of constructs usable for generating genetic intervals (see main text for details); RFP is driven by the *Sz. japonicus SJAG_04227* (*eis1*) promoter in pALo and by *Sz. octosporus SPOG_00147* (*pil2*) promoter in pALo181, CFP by the *Sz. octosporus SPOG_00147* (*pil2*) promoter in pALo & pALo182, and the yellow-shifted GFP* by the *Sz. cryophilus SOCG_04642* (*pil2*) promoter in pALo & pALo186.

### Monitoring meiosis chromosome segregation defects

Meiotic chromosome segregation defects can be monitored by markers inserted next to the centromere. Previously, this has been exploited in genetic screens by introducing bacterial operator repeats (*lacO* or *tetO*) close to centromeres in budding and fission yeast, to identify chromosome segregation mutants via the distribution of LacI- or TetR-GFP fusions binding to their respective operators, thus becoming visible as small foci (Straight et al. 1996; Michaelis et al. 1997; Nabeshima et al. 1998; Katis et al. 2004; Rabitsch et al. 2004; Gregan et al. 2005). Introducing spore-autonomously expressed fluorophore markers with different colors at the centromere (Figs. 1b, 2a) has the advantages of (I) enabling distinction of meiosis I and meiosis II segregation defects in a single assay (Fig. 2) rather than requiring homozygous and heterozygous setups of *lacO* or *tetO* repeats integrated close to a centromere, and (II) likely not interfering with chromosome behavior as strongly as *lacO* or *tetO* repeats (Fuchs et al. 2002; Sofueva et al. 2011). Fission yeast asci are ordered, due to the physical constraints of the zygotic cell size and shape microtubular spindles can orientate only along the longitudinal axis of the zygote, which means that the neighboring nuclei/spores in one half of the zygote are the sister products generated in meiosis II (Fig. 2b). This makes the evaluation of chromosome mis-segregation a comparatively straightforward undertaking in *Sz. pombe*.

**Figure 2.**
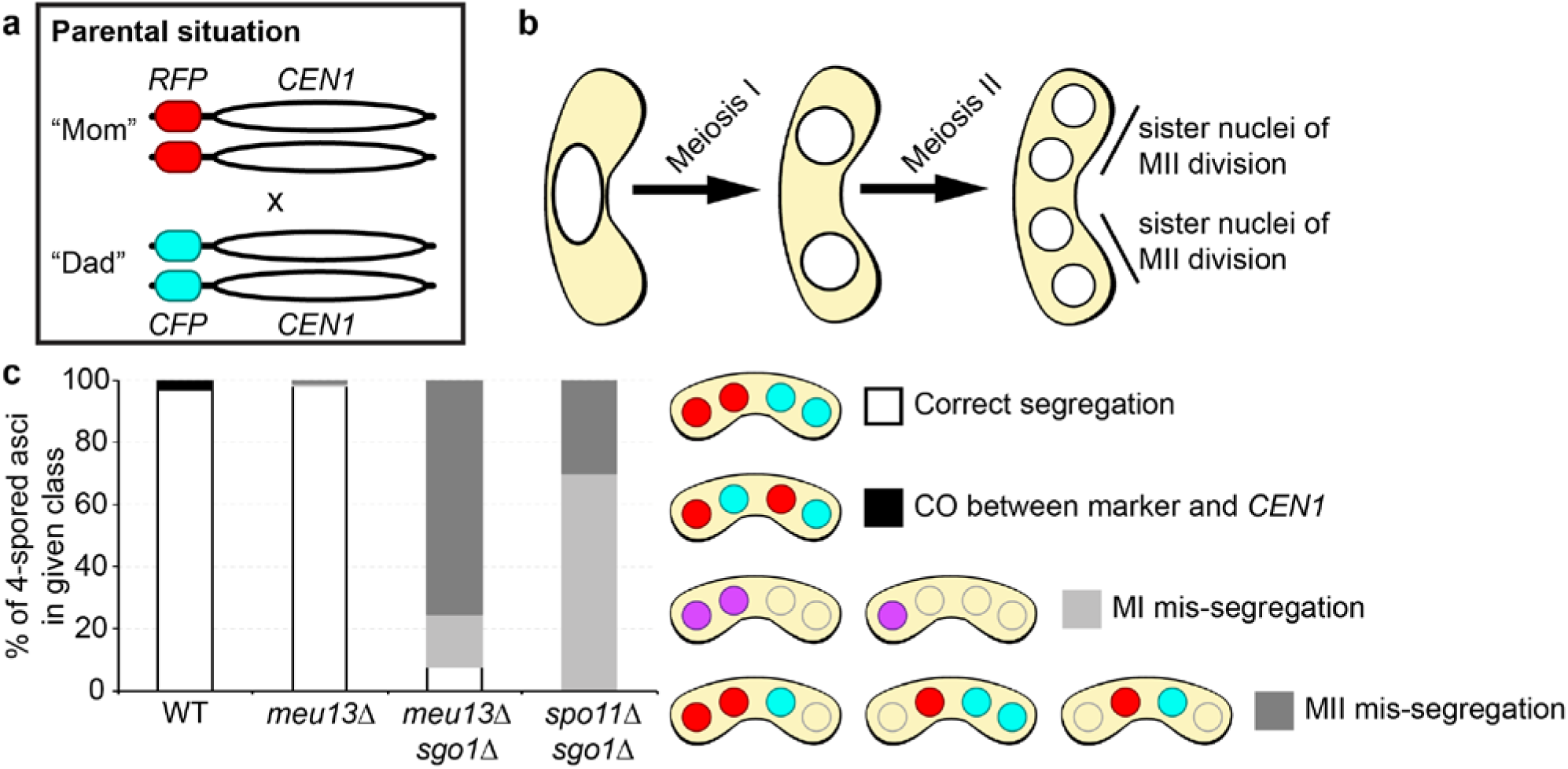
Chromosome segregation assay using spore-autonomous expression of fluorophores. (**a**) Schematic of assay, RFP and CFP are expressed from the *Sz. octosporus SPOG_00147* (*pil2*) promoter integrated at position 3,751, on chromosome I downstream of the *per1* (*SPAP7G5.06*) locus close to its centromere (*CEN1*). (**b**) Meiotic nuclear divisions generate an ordered tetrad with sister nuclei from meiosis II (MII) ending up next to one another. (**c**) Chromosome segregation phenotypes in 4-spored wild-type (WT; UoA585726×UoA585727, n = 274), *meu13*Δ (UoA585752×UoA585755, n = 101), *meu13*Δ *sgo1*Δ (UoA585756×UoA585759, n = 53), and *spo11*Δ *sgo1*Δ (UoA585760×UoA585763, n = asci) asci. A low frequency of crossover (CO) events (3.3%) between the fluorophore genes and *CEN1* have been observed in WT.

We tested the functionality of our assay with a set of mutants defective in meiotic recombination (*meu13*, *spo11*) and/or kinetochore function (*sgo1*) (Keeney et al. 1997; Nabeshima et al. 2001; Sharif et al. 2002; Rabitsch et al. 2004). For this we only evaluated 4-spore asci, and ignored asci with spore counts of 1, 2, or 3, to exclude incidences of clear nuclear division failures in meiosis I or II. As expected, in wild-type and *meu13*Δ crosses chromosome I is correctly segregated, in almost all cases (Fig. 2c). We did observe a low frequency (3.3%) of CO recombination between the fluorophore marker and the physical centromere in wild type, leading to red-cyan pairs of sister nuclei, rather than red-red and cyan-cyan pairs (Fig. 2c). In *meu13*Δ, which strongly reduces meiotic recombination (Nabeshima et al. 2001), no COs were observed, but incidences of chromosome mis-segregation could be recorded (Fig. 2c). As an obvious example for meiotic chromosome mis-segregation, we employed double mutants of *sgo1*Δ with *meu13*Δ or *spo11*Δ. A *sgo1*Δ single mutant does not produce a strong mis-segregation phenotyp (Rabitsch et al. 2004), but in combination with the absence of recombination factors a meiotic non-disjunction phenotype can be observed (Hirose et al. 2011). Indeed, massive chromosome segregation defects are obvious in asci of *meu13*Δ *sgo1*Δ and *spo11*Δ *sgo1*Δ double mutants (Fig. 2c). In *spo11*Δ *sgo1*Δ the percentage meiotic non-disjunction is slightly higher than in *meu13*Δ *sgo1*Δ, and there are also more meiosis I chromosome mis-segregation events in *spo11*Δ *sgo1*Δ. In *meu13*Δ chromosome segregation can presumably be supported to some degree, because a small number of chiasmata is still being produced, whereas in *spo11*Δ meiotic DSB formation is completely abrogated and thus no chiasmata are formed.

### Creating a genetic interval with fluorophore markers to assess meiotic recombination frequency

To explore whether fluorophore markers inserted at defined genomic sites on a single chromosome to create a genetic interval that can be used to determine meiotic recombination frequencies, we transformed constructs integrating on chromosome III forming a genetic interval of ~45kb around the *ade6* locus (Fig. 3a, see supplementary materials for details).

**Figure 3.**
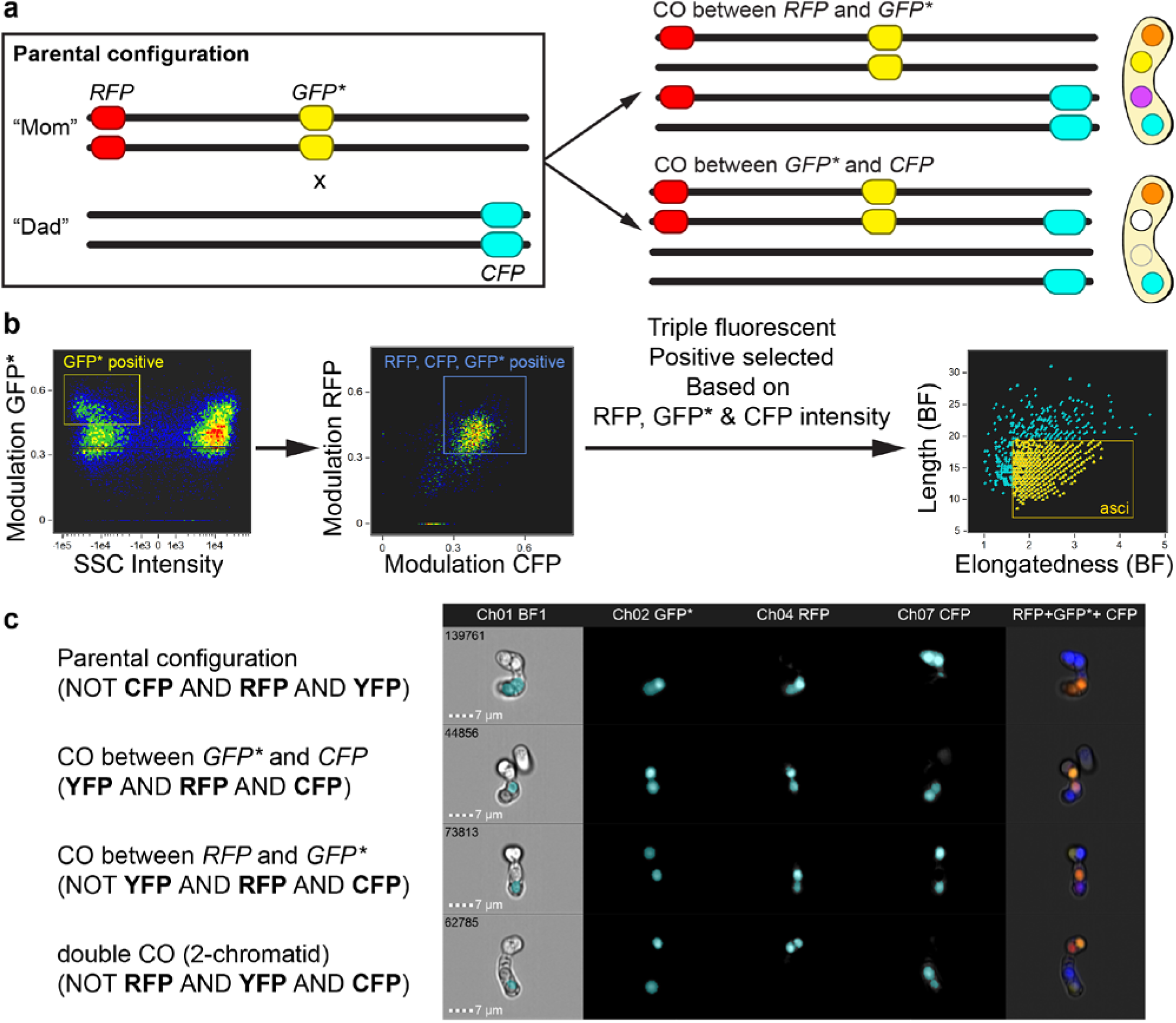
Genetic interval constructed with spore-autonomously expressed fluorescent markers can be analyzed by imaging flow cytometry. (**a**) Schematic of the genetic interval constructed: *RFP* expressed from the *Sz. japonicus SJAG_04227* (*eis1*) promoter together with a *ura4*+ marker is integrated on chromosome III at position 1,291,583, the same site as *ura4*+-*aim2* (Osman et al. 2003); *CFP* expressed from the *Sz. octosporus SPOG_00147* (*pil2*) promoter together with a *his3*+ marker is integrated on chromosome III at position 1,337,447, the same site as *his3*+-*aim* (Osman et al. 2003); *GFP*\* expression driven from the *Sz. cryophilus SOCG_04642* (*pil2*) promoter, the construct is integrated close to *ade6* at its endogenous locus. Only outcomes of single crossovers (COs) between the three markers are shown, double COs are rare (see Fig. S2 & Fig. S3 for double COs observed in this kind of assay). Please note, that order of spore colors is not fixed, but can rotate perpendicular to the meiotic spindle axis. (**b**) Outline of the workflow to identify asci based on particular cellular features on the Amnis ImageStreamX Mark II. Modulation measures the intensity range of an image normalized between and by calculating Modulation = (Max Pixel - Min Pixel)/(Max Pixel + Min Pixel). (**c**) Examples of ascus phenotypes from a cross of wild-type strains (UoA585694×UoA585676) as shown in (a); Boolean algebra mask equations used to discriminate between the different ascus types as presented in Ch BF1.

As this assay visualizes recombination events, we evaluated it using standard epi-fluorescence microscopy, and also tested whether single-cell imaging flow cytometry (Basiji 2016) could be exploited to perform high-throughput screens with the spore-autonomously expressed fluorophore recombination assay. We established a workflow on the Amnis ImageStreamX Mark II imaging flow cytometer to select for mature asci displaying fluorescence from a mixed population of cells in a standard cross (mature fluorescing asci, immature non-fluorescing asci, zygotes, vegetative cells), and subsequently applied customizations in the software to identify spore color phenotypes unique for the recombination outcomes we expected to occur in this assay (Fig. 3b,c).

Because the fluorophore markers were inserted at the same positions as the nutritional markers of an established recombination assay (Figs. 4a,b and S1a,b) (Osman et al. 2003; Lorenz et al. 2012, 2014), we could directly compare the outcomes of the different assays assessed by various methods. We used two slightly different recombination assays utilizing nutritional markers: one contained a point mutation at *ade6* (*ade6-704*, a T645A substitution mutation; Park et al. 2007), the other one a dominant drug resistance marker inserted at the 3’ end of *ade6* creating a partial deletion (*ade6-3’*Δ::*natMX6*). The latter was used to test whether a drastically different recombination frequency is caused by introducing a heterologous piece of DNA into the genetic interval. The *natMX6* cassette is ~1.25kb in size and removes 842bp (426bp of which are *ade6* coding sequence), in comparison the *GFP*\* cassette is ~2.1kb in size and inserted just downstream of the *ade6* open reading frame (see supplementary materials for details of strain construction).

**Figure 4.**
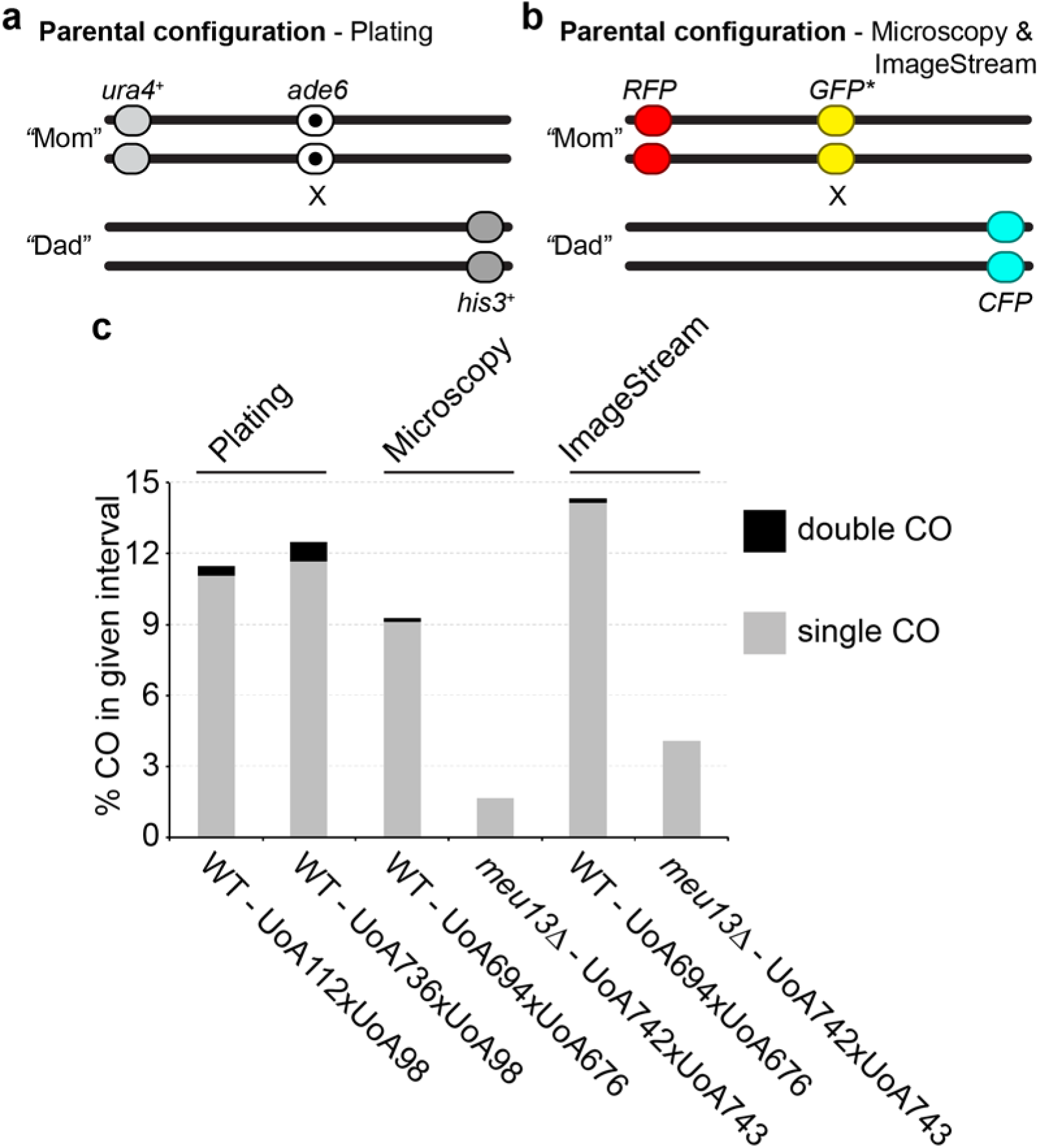
Comparison of genetic intervals generated by nutritional markers and spore-autonomously expressed fluorescent markers. (**a**) Schematic of genetic recombination assay using nutritional markers and plating of colonies. In UoA112 the *ade6* marker is a point mutation *(ade6-704*) without hotspot activity, in UoA585 it is a partial deletion of *ade6* by integrating a *natMX6* cassette (*ade6-3’*Δ) examples of main constructs. The flanking markers *ura4*+ and *his3*+ are the artificially introduced markers (aim) *ura4*+-*aim2* and *his3*+-*aim*, which have been previously described (Osman et al. 2003). (**b**) Schematic of spore-autonomously expressed fluorophore recombination assay (see also Fig. 3a), the *RFP* gene is at the same position as *ura4*+-*aim2* in (a), the *CFP* gene at the same position as *his3*+-*aim* in (a), and the *GFP*\* gene is inserted downstream of *ade6*+. (**c**) Results from recombination assays in (a) and (b): crossover (CO) recombinant frequencies were determined in wild-type (WT) crosses by random spore analysis for the plating assay (a), using data from n = independent crosses with progeny each. CO recombinant frequencies were determined in WT and *meu13*Δ crosses either by counting manually on an epi-fluorescence microscope (UoA585694×UoA585676n = 356asci, UoA585742×UoA585743n = 305asci) or by high-throughput single cell assessment on an imaging flow cytometer (ImageStream) (UoA585694×UoA585676n = 916asci, UoA585742×UoA585743n = 370asci). Please note, that ImageStream can only identify one out of two double CO classes.

Despite all these differences between the genetic markers, the recombination frequencies within the genetic intervals were remarkably similar (Figs. 4c and S1c). The genetic intervals with the nutritional markers produced 11.88% (*ade6-704*) and 13.33% (*ade6-3*’Δ) COs, respectively (Figs. 4c, Table S4). The interval with the fluorophore markers measured 9.41% COs on the epi-fluorescence microscope and 14.57% COs on the imaging flow cytometer (Fig. 4c, Table S5). The results were comparable, when the *ade6*- or *GFP*\*-markers were initially linked with *his3*+-*aim* or *CFP*, respectively (10.63% CO for ade6-704, 8.33% CO for *ade6-3’*Δ, 7.68% CO for fluorophore markers evaluated by epi-fluorescence microscopy; Fig. S1, Tables S4 and S5). In all types of assays we could also detect a few rare double CO events (Figs. 4 and S1, Tables S4 and S5). Because asci can be evaluated as an ordered tetrad in the fluorophore-based assay (Figs. 2b and 3a), information about the involvement of 2, 3, or all 4 chromatids in the double CO can be extracted. Within the 4 double CO events over the two slightly different genetic intervals evaluated on the epi-fluorescence microscope (Figs. 4b and S1b), examples for participation of 2, 3, or 4 chromatids could be found (Figs. S2 and S3). The observed frequency of double CO in any of the genetic assaysis equal with or slightly higher than expected from the frequency in neighboring intervals (Table SX), in line with Sz. pombe not displaying CO interference (Munz 1994).

In a *meu13* mutant meiotic intra- and intergenic recombination is strongly decreased (Nabeshima et al. 2001). When running the fluorophore-based assay in a *meu13*Δ background, as expected, a 3.6- to 5.7-fold reduction in CO formation could be observed (Fig. 4c). No double COs were detected in the *meu13*Δ crosses. This demonstrates that in *Sz. pombe* a genetic interval consisting of spore-autonomously expressed fluorescent markers behaves very similarly to a genetic interval built from nutritional markers.

### Conclusion

Here, we established assays employing spore-autonomously expressed fluorescent proteins to determine meiotic chromosome mis-segregation and meiotic recombination frequencies in the fission yeast, *Sz. pombe*. We generated a series of plasmids containing selectable markers (*ura4*+, *his3*+, and *arg3*+) in addition to the spore-specific fluorophores (Fig. 1, Table S1); this makes the whole system portable enabling the creation of genetic intervals at virtually any position within the *Sz. pombe* genome. Ectopic spore-autonomous promoters from *Sz. japonicus* work in *Sz. pombe*, this raises the possibility that expression from this type of regulatory elements is conserved, and could be used to develop a similar system in *Sz. japonicus*. This is of interest, because *Sz. japonicus* produces 8-spored asci (an additional mitosis following the two meiotic divisions) (Klar 2013) enabling an even better resolution of genetic events. We validated our system by comparison to an established recombination assay (Osman et al. 2003; Lorenz et al. 2012, 2014) utilizing nutritional markers (Fig. 4), and demonstrated that imaging flow cytometry can be used to run genetic high-throughput screens for recombination phenotypes (Figs. 3 and 4). Due to its portability and advantages over existing assays our fluorophore-based system represents a novel addition to the ever growing genetic toolkit for probing the cell biology of fission yeast.

## Acknowledgements

We are grateful to Scott Keeney, Franz Klein, Jürg Kohli, Josef Loidl, Kim Nasmyth, Ken E. Sawin, Gerald R. Smith, Takashi Toda, Matthew C. Whitby, and the National BioResource Project (NBRP) Japan for providing strains and/or plasmids, and to M.N. Asogwa, A. Bebes, and L. Duncan for technical assistance. Microscopy was performed at the University of Aberdeen Microscopy & Histology facility (Kevin Mackenzie). This work was supported by a Carnegie Trust for the Universities of Scotland Research Incentive Grant (No. 70021), and the University of Aberdeen (College of Life Sciences and Medicine Start-up grant).

## Supplementary Materials

**Figure S1.**
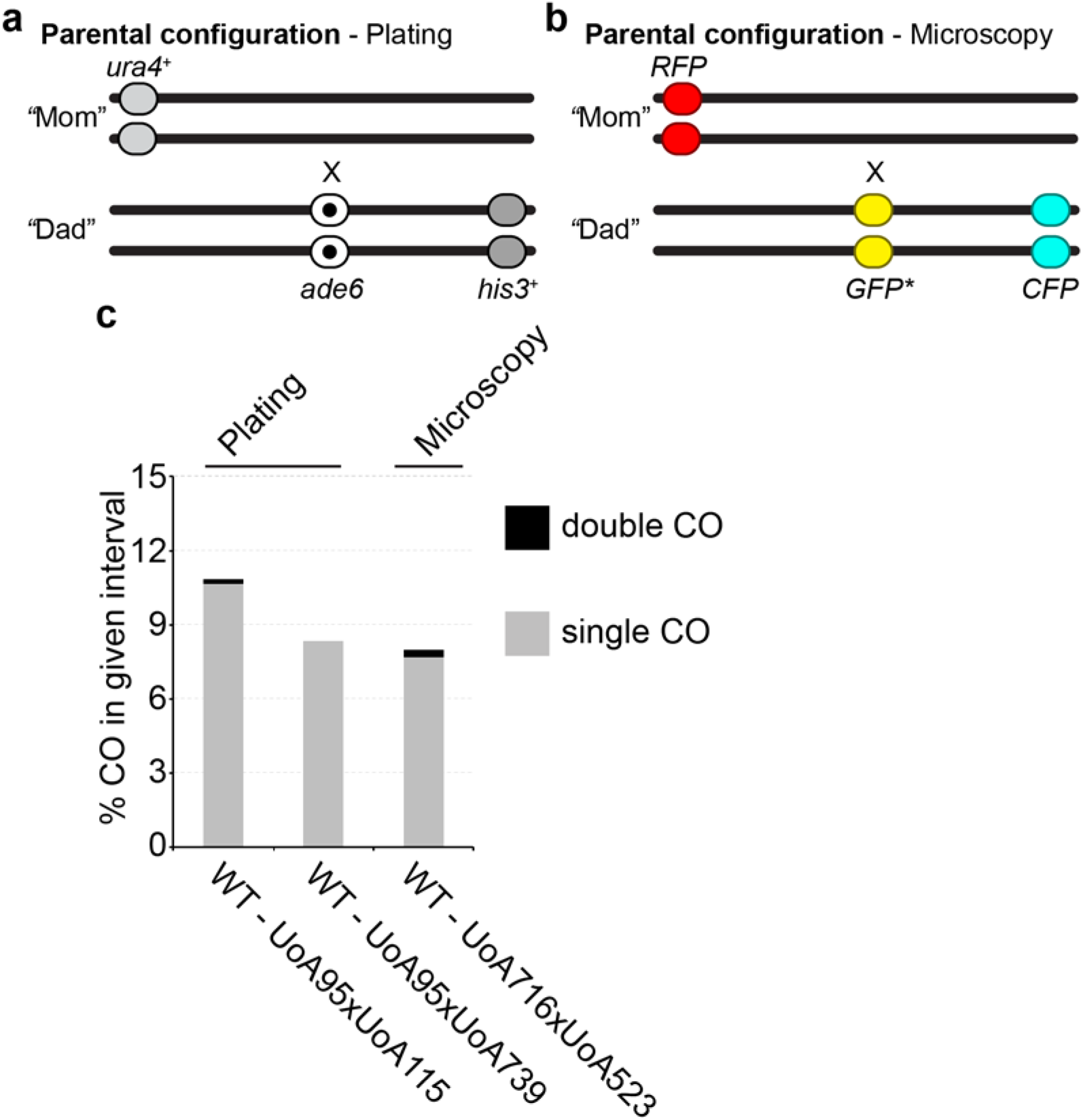
Comparison of genetic intervals generated by nutritional markers and spore-autonomously expressed fluorescent markers. (**a**) Schematic of genetic recombination assay using nutritional markers and plating of colonies. In UoA115 the *ade6* marker is a point mutation *(ade6-704*) without hotspot activity, in UoA585 it is a partial deletion of *ade6* by integrating a *natMX6* cassette (*ade6-3’*Δ) examples of main constructs. The flanking markers *ura4*+ and *his3*+ are the artificially introduced markers (aim) *ura4*+-*aim2* and *his3*+-*aim*, which have been previously described (Osman *et al*. 2003). (**b**) Schematic of spore-autonomously expressed fluorophore recombination assay, the *RFP* gene is at the same position as *ura4*+-*aim2* in (a), the *CFP* gene at the same position as *his3*+-*aim* in (a), and the *GFP*\* gene is inserted downstream of *ade6*+. (**c**) Results from recombination assays in (a) and (b): crossover (CO) recombinant frequencies were determined in wild-type (WT) crosses by random spore analysis for the plating assay (a), using data from n = independent crosses with progeny each. CO recombinant frequencies were determined in WT and *meu13*Δ crosses either by counting manually on an epi-fluorescence microscope (UoA585716=UoA585523 n = 508 asci).

**Figure S2.**
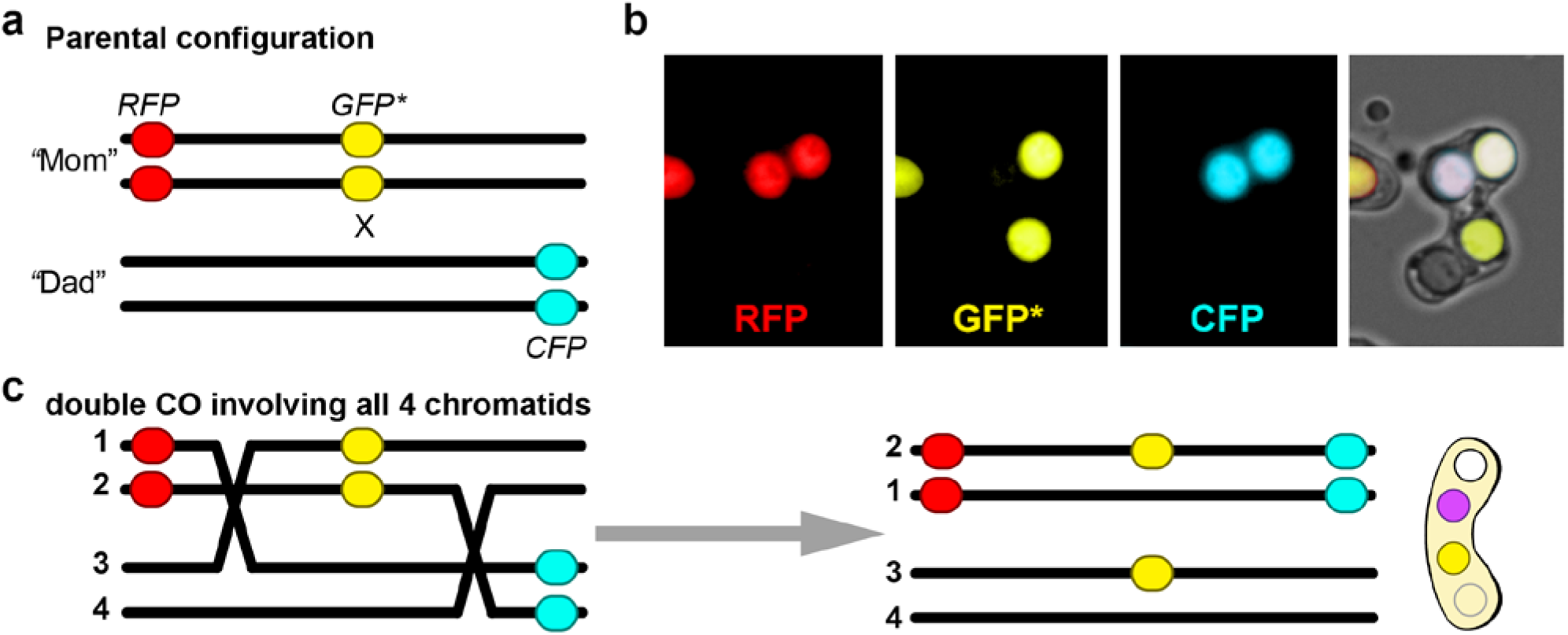
Double crossover (CO) observed in genetic interval with spore-autonomously expressed fluorescent markers. (**a**) Schematic of spore-autonomously expressed fluorophore recombination assay for cross UoA585694×UoA585 (see also Figs. 3 & 4). (**b**) The only double CO event observed among asci; RFP, GFP*, and CFP fluorescence channels are shown separately and as a merged image. (**c**) Most parsimonious explanation for this ascus phenotype is a double CO involving all chromatids (numbered on the left); possible alternative interpretations would require a third CO.

**Figure S3.**
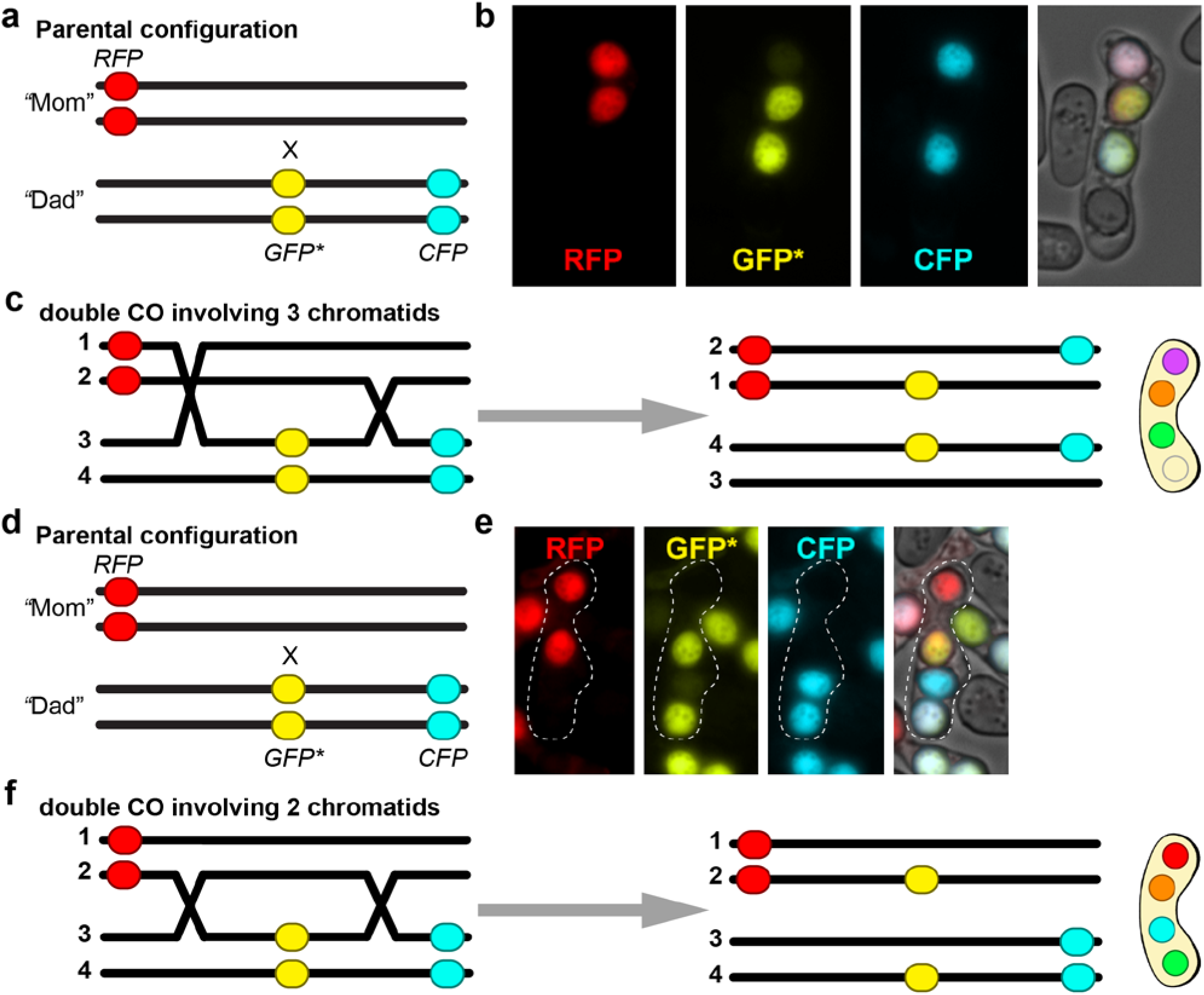
Double crossover (CO) observed in genetic interval with spore-autonomously expressed fluorescent markers. (**a**) Schematic of spore-autonomously expressed fluorophore recombination assay for cross UoA585716×UoA585 (see also Fig. S1). (**b**) Two double CO events observed among asci; RFP, GFP*, and CFP fluorescence channels are shown separately and as a merged image (one of the two asci is given as example). (**c**) Most parsimonious explanation for this ascus phenotype is a double CO involving chromatids (chromatids are numbered on the left). (**d**) Schematic of spore-autonomously expressed fluorophore recombination assay for cross UoA585716×UoA585523 (see also Fig. S1). (**e**) One double CO events observed among asci; RFP, GFP*, and CFP fluorescence channels are shown separately and as a merged image. (**f**) Most parsimonious explanation for this ascus phenotype is a double CO involving chromatids only (chromatids are numbered on the left).

**Supplementary Table S1.**
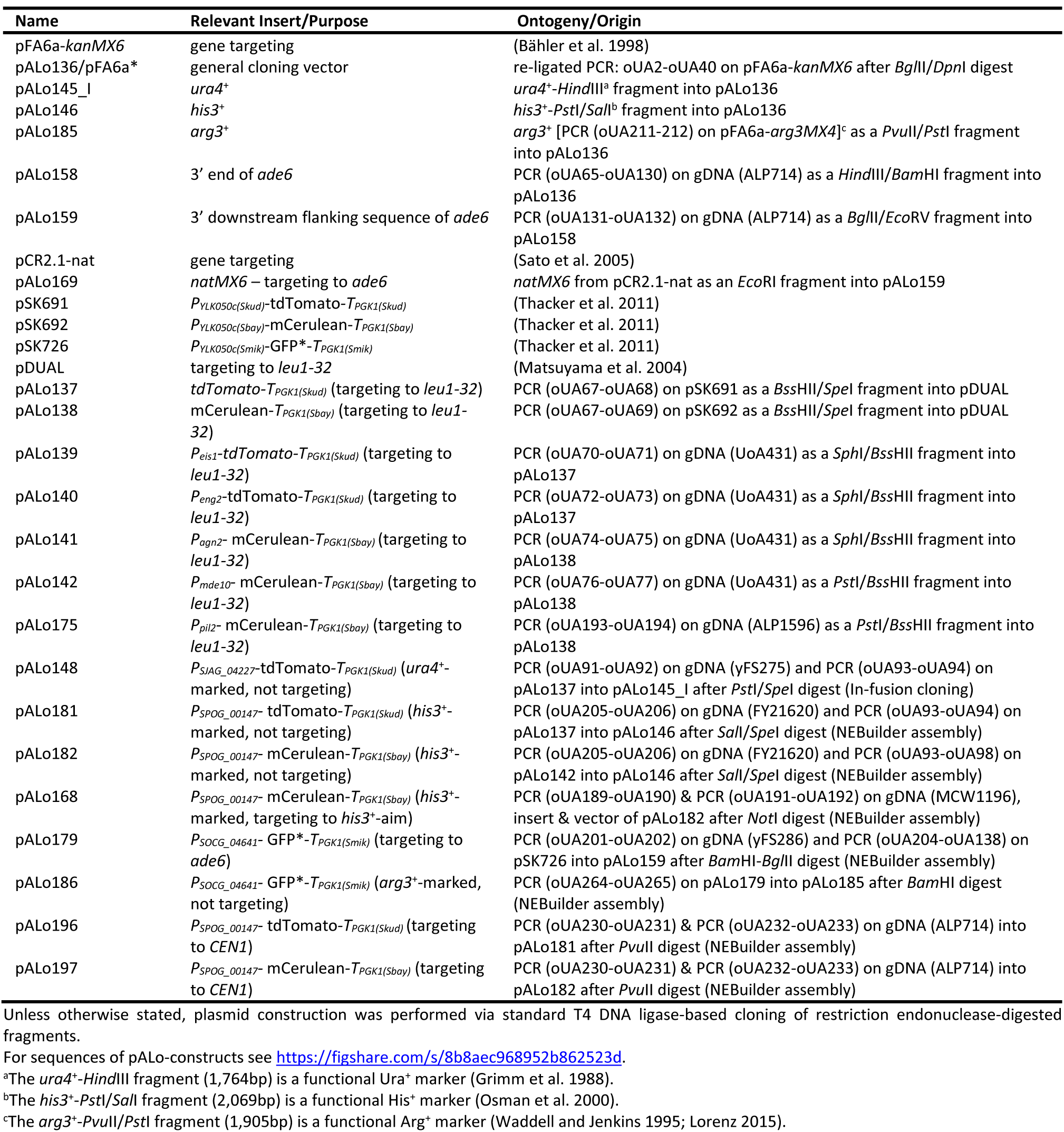
Plasmid list.

**Supplementary Table S2.**
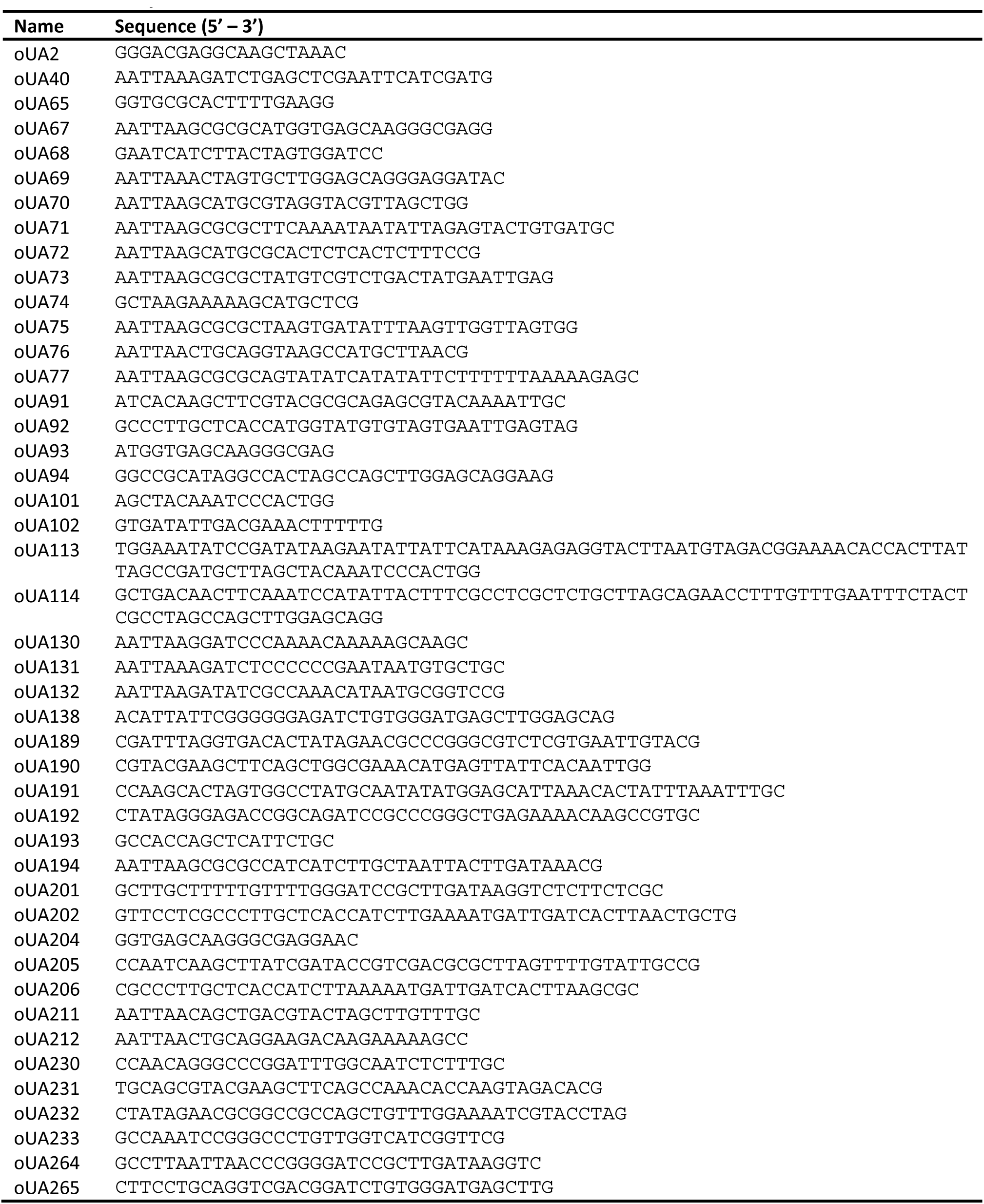
List of Oligonucleotides.

**Supplementary Table S3.**
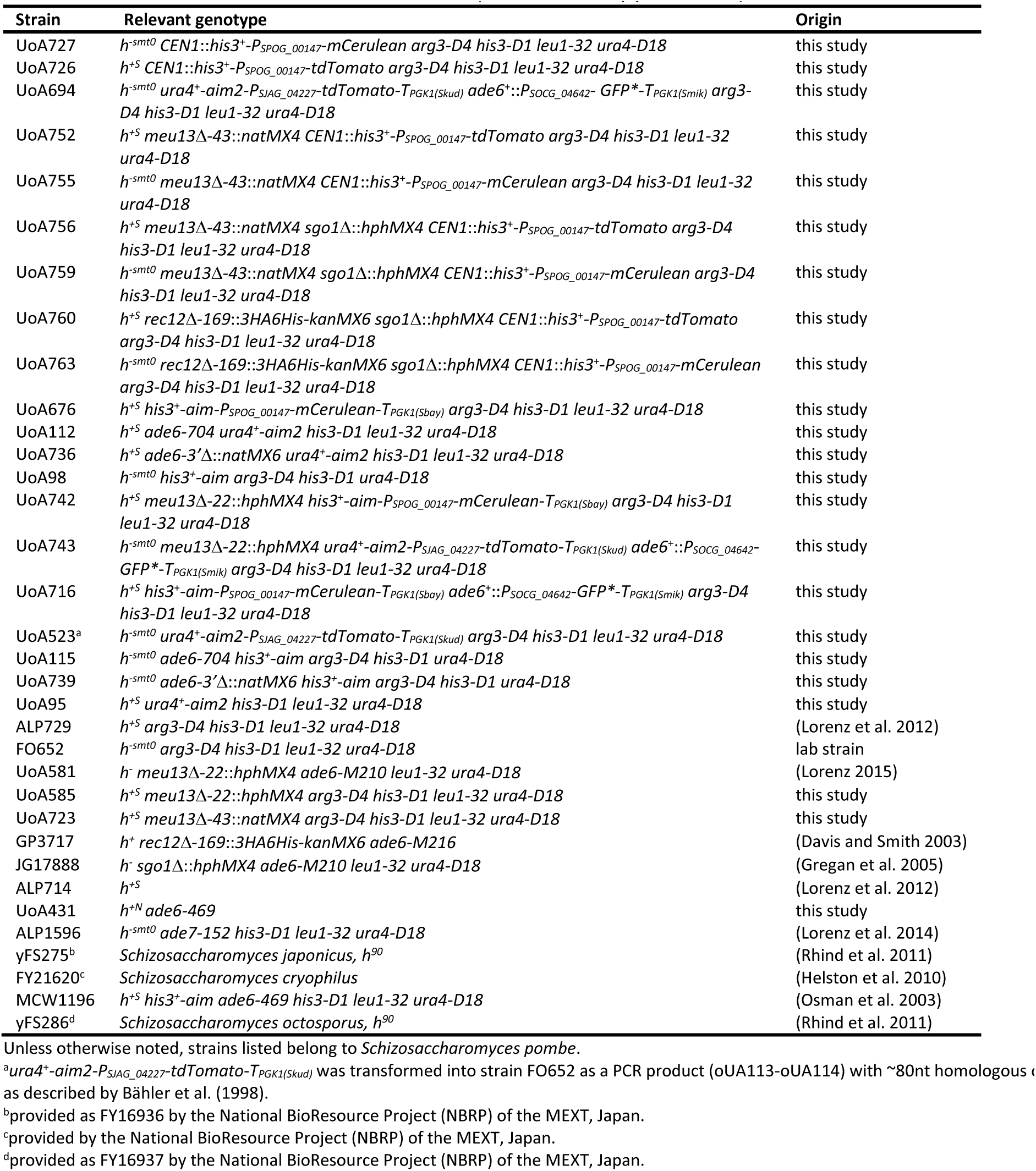
Yeast strain list (in order of appearance)

**Supplementary Table S4.** Recombination frequency in the interval *ura4*+-*aim2* – *ade6-704*/*ade6-3’*Δ – *his3*+-*aim* by random spore analysis.

| cross | % spore viability [±S.D.] | % total CO [±S.D.] | % CO *ura4-ade6* [±S.D.] | % CO *ade6-his3* [±S.D.] | % double CO [±S.D.] | % double CO (exp.) |
| --- | --- | --- | --- | --- | --- | --- |
| UoA112×UoA98 | 74.7 [±7.84] | 11.88 [±4.1] | 6.88 [±3.13] | 5.0 [±1.08] | 0.42 [±0.72] | 0.34 |
| UoA736×UoA98 | 74.6 [±5.59] | 13.33 [±1.3] | 5.83 [±0.96] | 7.5 [±1.25] | 0.83 [±0.36] | 0.44 |
| UoA95×UoA115 | 87.6 [±2.91] | 11.04 [±2.95] | 6.67 [±2.95] | 4.38 [±0.0] | 0.21 [±0.36] | 0.74 |
| UoA95×UoA739 | 76.7 [±15.1] | 8.33 [±3.08] | 6.04 [±1.8] | 2.29 [±1.3] | 0.0 | 0.14 |

Crossover (CO) frequency expressed as (number of recombinant progeny)/(total viable colonies) in %.

**Supplementary Table S5.** Recombination frequency in the interval *RFP* – *GFP*\* – *CFP*.

| cross | % total CO | % CO *RFP-GFP\** | % CO *GFP\*-CFP* | % double CO | % double CO (exp.) |
| --- | --- | --- | --- | --- | --- |
| UoA716×UoA523 | 8.29 | 1.97 | 6.3 | 0.295 | 0.124 |
| UoA694×UoA676 | 9.41 | 1.55 | 7.87 | 0.14 | 0.121 |
| UoA742×UoA743 | 1.63 | 0.33 | 1.31 | 0.00 | 0.0043 |

Crossover (CO) frequency expressed as (2× number of recombinant asci)/(4× total number of asci) in %.

